# Predicting evolutionary change at the DNA level in a natural Mimulus population

**DOI:** 10.1101/2020.06.23.166736

**Authors:** Patrick J. Monnahan, Jack Colicchio, Lila Fishman, Stuart J. Macdonald, John K. Kelly

## Abstract

Evolution by natural selection occurs when the frequencies of genetic variants change because individuals differ in Darwinian fitness components such as survival or reproductive success. Differential fitness has been demonstrated in field studies of many organisms, but our ability to quantitatively predict allele frequency changes from fitness measurements remains unclear. Here, we characterize natural selection on millions of Single Nucleotide Polymorphisms (SNPs) across the genome of the annual plant *Mimulus guttatus*. We use fitness estimates to calibrate population genetic models that effectively predict observed allele frequency changes into the next generation. Hundreds of SNPs experienced “male selection” in 2013 with one allele at each SNP elevated in frequency among successful male gametes relative to the entire population of adults. In the following generation, allele frequencies at these SNPs consistently shifted in the predicted direction. A second year of study revealed that SNPs had effects on both viability and reproductive success with pervasive trade-offs between fitness components. SNPs favored by male selection were, on average, detrimental to survival. These trade-offs (antagonistic pleiotropy and temporal fluctuations in fitness) may be essential to the long-term maintenance of alleles undergoing substantial changes from generation to generation. Despite the challenges of measuring selection in the wild, the strong correlation between predicted and observed allele frequency changes suggests that population genetic models have a much greater role to play in forward-time prediction of evolutionary change.

**Author summary:** For the last 100 years, population geneticists have been deriving equations for Δp, the change in allele frequency owing to mutation, selection, migration, and genetic drift. Seldom are these equations used directly, to match a prediction for Δp to an observation of Δp. Here, we apply genomic sequencing technologies to samples from natural populations, obtaining millions of observations of Δp. We estimate natural selection on SNPs in a natural population of yellow monkeyflowers and find extensive evidence for selection through differential male success. We use the SNP-specific fitness estimates to calibrate a population genetic model that predicts observed Δp into the next generation. We find that when male selection favored one nucleotide at a SNP, that nucleotide increased in frequency in the next generation. Since neither observed nor predicted Δp are generally large in magnitude, we developed a novel method called “haplotype matching” to improve prediction accuracy. The method leverages intensive whole genome sequencing of a reference panel (187 individuals) to infer sequence-specific selection in thousands of field individuals sequenced at much lower coverage. This method proved essential to accurately predicting Δp in this experiment and further development may facilitate population genetic prediction more generally.

## Introduction

Natural selection is routinely strong enough to measure within natural populations. Classic experiments on conspicuous polymorphisms were the first to demonstrate fitness differences among genotypes [1, 2]. Field experiments later demonstrated selection on allozymes [3] and structural variants such as inversions [4–6], but quantitative trait locus (QTL) mapping greatly expanded the set of loci amenable to direct study [7]. The link that QTLs provide to phenotype can enable a “mechanistic” understanding of selection, allowing us to describe the processes that maintain polymorphism (e.g. antagonistic pleiotropy [4, 8], frequency dependent selection [9] or gametic/zygotic fitness trade-offs [10]), and the environmental drivers of selection (e.g. differential predation [11]). In aggregate, these single-locus studies have provided great insight on the contribution of major loci to the standing variance in fitness within natural populations.

Genome-wide surveys of natural populations deliver a comprehensive view of selection. An important question is how many loci across the genome experience selection in a typical generation. Sequencing of natural populations sampled through time suggests that the strong selection documented in single locus studies can occur at hundreds of polymorphisms simultaneously [12, 13]. In *Drosophila melanogaster*, large amplitude fluctuations in allele frequency occur seasonally and can be directly related to weather conditions [14]. The magnitude and consistency of changes, as well as the environmental correlation, clearly imply that selection (and not genetic drift) is causal. The temporal sampling method employed for *D. melanogaster* should be expanded to other systems in the future, but some questions require individual level genome sequence data. For instance, are fitness differences caused mainly by differences in viability or fertility or mating success? Experiments predicting individual fitness from individual genomes have been conducted in a variety of organisms using both “common gardens,” where sequenced individuals are transplanted into natural settings [15–18], as well as monitoring of native individuals *in situ* [19–21]. These studies yield varying results on the importance of different selection components, but in aggregate, suggest that selection is a pervasive force on ecological time scales.

Here, we measure genome-wide selection and allele frequency change in a field study of *Mimulus guttatus*; a plant species in which the various methods described above have been applied extensively within a single natural population at Iron Mountain (IM). We have demonstrated strong fitness effects of segregating inversions by genotyping IM plants that were also scored for fecundity [22, 23]. Transplant experiments using QTL constructs for ecologically important traits have confirmed that conflicting selection pressures are key to the maintenance of variation [24, 25]. QTL alleles that increase plant size at reproduction nearly always delay flowering, which generates antagonistic pleiotropy between survival and fecundity. These single-locus experiments (QTLs and inversions) have been corroborated by Genome Wide Association (GWA) of traits and fitness components in IM [17]. ‘Big/slow’ alleles that delay progression to flowering, but increase flower size, segregate at many loci across the genome. They tend to be less frequent than their ‘small/fast’ alternatives within IM [17, 26], which is consistent with many years of field monitoring indicating that viability selection generally favors small/fast alleles [24, 25, 27]. However, the GWA also demonstrated temporal fluctuation in the net balance of fitness components [17] suggesting that year-to-year changes in water availability are key to the maintenance of variation.

The focus of this paper is prediction: Can we characterize selection at the SNP level accurately enough to predict allele frequency change into the next generation? Prospective (forward-time) prediction of evolutionary change from measurements of selection is a primary goal of quantitative genetics [28–32], but has long been considered beyond the scope of population genetics [33]. In quantitative genetics, estimates of phenotypic selection (differentials or gradients) can be combined with estimates of inheritance (heritability or genetic (co)variance) to predict the change in mean phenotypes [34, 35]. Prediction accuracy can be improved by directly relating the loci affecting a trait to fitness, using either the secondary theorem of selection [36, 37] or via genomic selection methods [38]. The scope of quantitative genetics is broad, but its enduring relevance to both agriculture [39, 40] and evolutionary biology [29] owes importantly to its capacity for prospective prediction. It is an open question whether selection on SNPs strong enough to predict Δ*p*, the change in allele frequency, in a manner analogous to 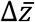, the change in mean phenotype.

To estimate selection on SNPs, we collected paired-end sequence reads from reduced representation [41] sequencing libraries of 1936 experimental plants (field individuals and progeny). We called variants within reads and aligned them to 187 full genome sequences previously obtained from the IM population [17]. This alignment is the basis for the “haplotype matching” technique of genotype inference. Below, we describe this technique and then provide a proof-of-concept application to data from the *Drosophila* Synthetic Population Resource (DSPR) [42] where haplotype inheritance is known. We then apply haplotype matching to derive genotype probabilities for SNPs within 15,360 genic regions of experimental plants. These likelihoods are inputs to the selection component models that predict allele frequency change [19, 43]. Male selection is measured by synthesizing maternal and progeny sequencing to infer the (unseen) male siring fitness component. We show that male selection in 2013 predicts observed changes in allele frequency into the next generation; the latter estimated from a distinct sampling of plants in 2014.

**Figure 1.**
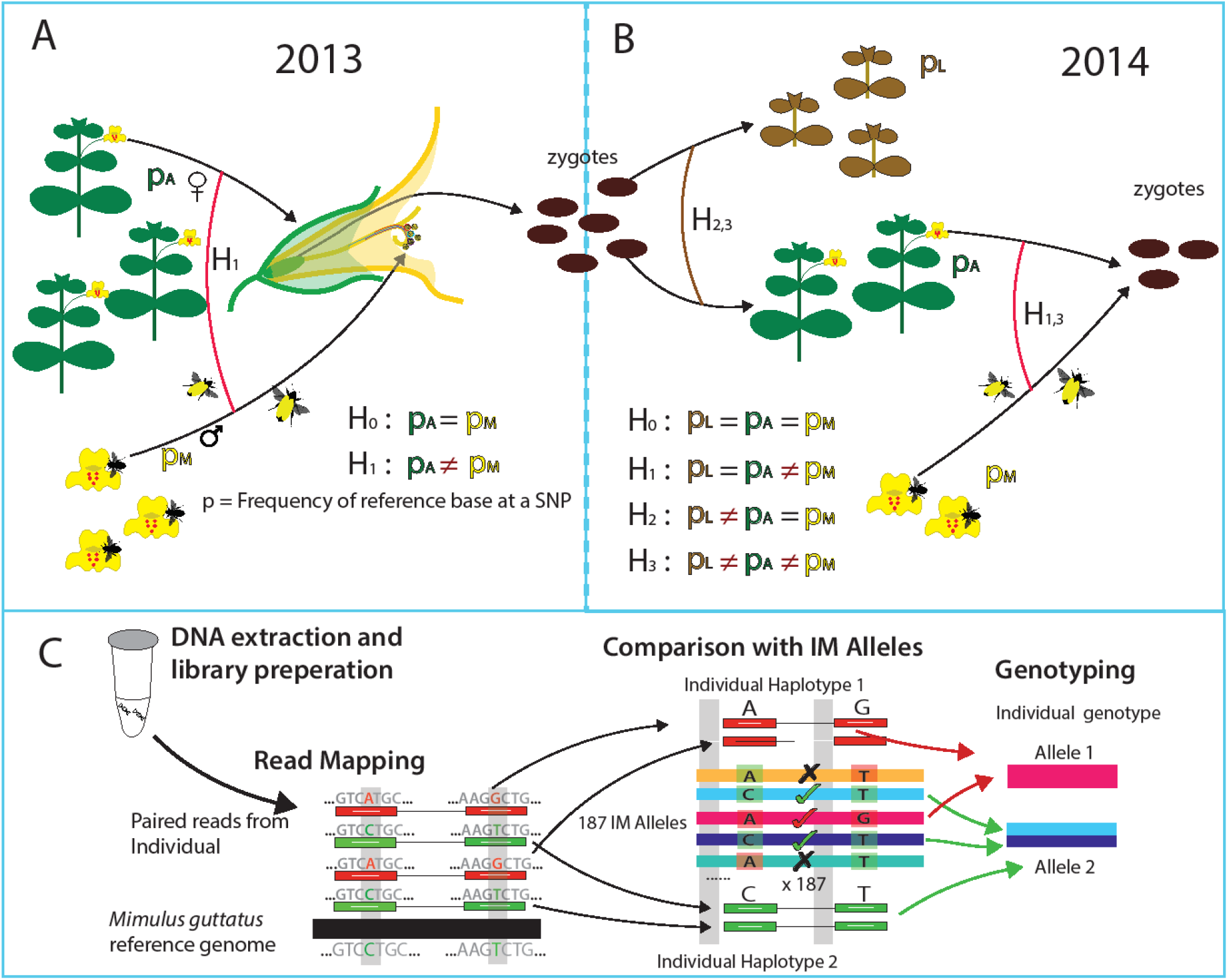
The parameters of alternative selection models are depicted for the (A) 2013 and (B) 2014 data. Hypothesis tests are expressed in terms of parameter constraints where p indicates reference base frequency: p_A_ for reproductive adults, p_M_ for successful male gametes, and p_L_ plants for plants that fail to reproduce. H_0_ is the full neutral model. Male selection is tested by contrast of H_1_ to H_0_ in 2013 and H_3_ to H_1_ in 2014. Viability selection is tested by contrast of H_3_ to H_2_. (C) After DNA sequencing, read-pairs are mapped to the *M. guttatus* reference genome. The haplotype matching method (read-pairs to genic-haplotypes) is illustrated for a simple case with read-pairs mapping to single location. Read-pairs impose a probabilistic ‘process of elimination’ on reference line sequences as putative ancestors: √ indicates consistency and “X” inconsistency.

## Results and Discussion

*Mimulus guttatus* (syn. *Erythranthe guttata*) is a hermaphroditic species that can experience selection prior to flowering, via differential viability, and subsequent to flowering through both male and female function. In the first year of our study (Fig 1A: 2013), we sampled plants that successfully flowered (adults), genotyping them as well as a random collection of their progeny. Given the maternal genotype, we can statistically identify her allelic contribution to offspring and distinguish allele frequency among all adults (*p_A_*) from that in the population of *successful* male gametes (*p_M_*). The *p_A_/p_M_* test evaluates whether these frequencies are different and thus identifies selection through differential male success. “Male selection” integrates a number of distinct selective mechanisms [19] including simple differences in fecundity (which may be equivalent between male and female function), sexual selection through differential siring [44] and pollen competition [45].

To test the predicted changes caused by male selection in 2013, we sampled plants from the next generation (Fig. 1B: 2014). We used MSG-RADseq [41] reduced representation sequencing to genotype three distinct cohorts: individuals that germinated but failed to reproduce (allele frequency *p_L_*), individuals that successfully flowered and produced fruit (allele frequency *p_A_*), and a random sample of progeny from reproductive individuals (used to estimate *p_M_*). We performed statistical contrasts between cohorts, asking whether allele frequency differs using likelihood based selection component models [43, 46–48] generalized to accommodate uncertain genotype calls [19]. Selection is indicated when a model that allows allele frequencies to differ between cohorts, e.g. *p_A_* ≠ *p_M_*, has a much higher likelihood than a constrained model, e.g. *p_A_* = *p_M_* (see METHODS section D).

We derived SNP allele frequency estimates using a two-stage genotyping strategy (Fig 1C). Read-pairs are initially matched to the set of ‘genic haplotypes’ present in IM. Sequence variation is very high in *M. guttatus* [49] and it is difficult to effectively call variants outside genic regions. We thus established “gene sets” as loci. A set is either a single gene or a collection of closely linked (within 100bp) and/or overlapping genes (Supplemental Table S1). The genic haplotypes are the sequences for this locus among the reference panel genomes (detailed procedures in Supplemental Appendix B). With 187 distinct haplotypes, there are 17,578 distinct genic-genotypes. However, most gene sets have fewer than 187 because some IM lines are identical within a gene set (the median number of distinct genic-haplotypes is 100, Supplemental Table S1).

We treat the genic haplotypes as the sequences present in the natural population (Fig 1C). Let *U*_[*plantID*],*i,j*_ denote the likelihood for the full collection of read-pairs from a plant given that its diploid genic-genotype is [*i*,*j*], where *i* and *j* index genic haplotypes. For an outbred plant,

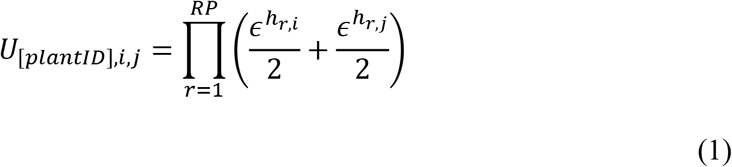

where RP is the number of read-pairs mapped in this gene set, *h_r,i_* is the number of sequence mismatches between read-pair *r* and genic haplotype *i*, and *ϵ* is the mismatch probability. *ϵ* aggregates the various events (sequencing error, alignment error, etc) that could create an apparent sequence difference even if the read-pair and haplotype are the same. *U* relates the RADseq data collected from field plants to the tests for selection.

A potential difficulty with haplotype matching is that the sequence of a field plant may not match any of our genic haplotypes; an error that could reduce our ability to detect selection. It is straightforward to test whether individual read-pairs are consistent with the genic haplotypes. Across the 99 million read-pairs in the final RADseq dataset (field plants from both years), the median number of SNPs per read-pair is 6. About 20% of read-pairs overlap 10 or more SNPs (Supplemental Table S2, Supplemental Figure S1). Across all read-pairs, less than 0.2% failed to perfectly match at least one genic haplotype. Of course, the full collection of read-pairs from a plant can still be inconsistent with any pair of genic haplotypes (even if all individual read-pairs map perfectly). This occurs, but very infrequently. In these cases, the genotype is treated as unknown. The consequences of incomplete sampling of the reference panel are explored in a companion paper [50].

Given consistency, the question becomes how precisely low-level sequencing can identify the genotype of field plants. As expected, the number of possible genic genotypes for a plant declines as the number of read-pairs mapped to gene set increases (Fig 2A). With low but reasonable coverage (10-20 read-pairs over an entire gene), the collection of compatible genic-genotypes is greatly reduced (on average to ≈5% of the total). Oftentimes, we identify one parental genic-haplotype definitively, but the other is consistent with multiple sequences from the reference set (illustrated by Fig 1C). The aggregation of evidence across numerous read-pair loci (mapping to different parts of gene) is usually needed to identify specific genic-haplotypes. While zeroing in on 5% of diploid genic-genotypes is still hundreds of possibilities, these possibilities often strongly “agree” about the genotype at particular SNPs – all or nearly all genic-genotypes have the same genotype at that SNP. SNP specific inference can be quite strong even with moderate coverage. Plants with low sequencing coverage often have few or no read-pairs, particularly in smaller gene sets. In isolation, inference for such plants would be weak. Here, inference can become much stronger with information from relatives (the maternal plant, siblings, or offspring). Importantly, we never truncate probabilities to produce “hard calls” for SNPs. Uncertainty is propagated through the entire analysis and thus properly integrated in testing. The selection analyses cycle through all SNPs within a gene set, considering each as a potential effector of fitness.

### A test of haplotype matching using Drosophila melanogaster

With the Mimulus data, we do not know the true genic-genotype of field plants and thus cannot compare inferred to known. For this reason, we applied our pipeline to a *Drosophila melanogaster* population where genic-genotypes are known with high confidence. The *Drosophila* Synthetic Population Resource (DSPR) consists of two multiparental, advanced generation intercross Recombinant Inbred Line (RIL) populations, each initiated from eight inbred founder strains [42, 51]. The founder strains have been fully sequenced and represent the reference panel in the current context. The RILs (comparable to Mimulus field plants) were genotyped and we know the founder strain that contributed the allele at each gene of each RIL. Some regions in some RILs are not genotyped with certainty, but we exclude these from our analyses.

**Figure 2.**
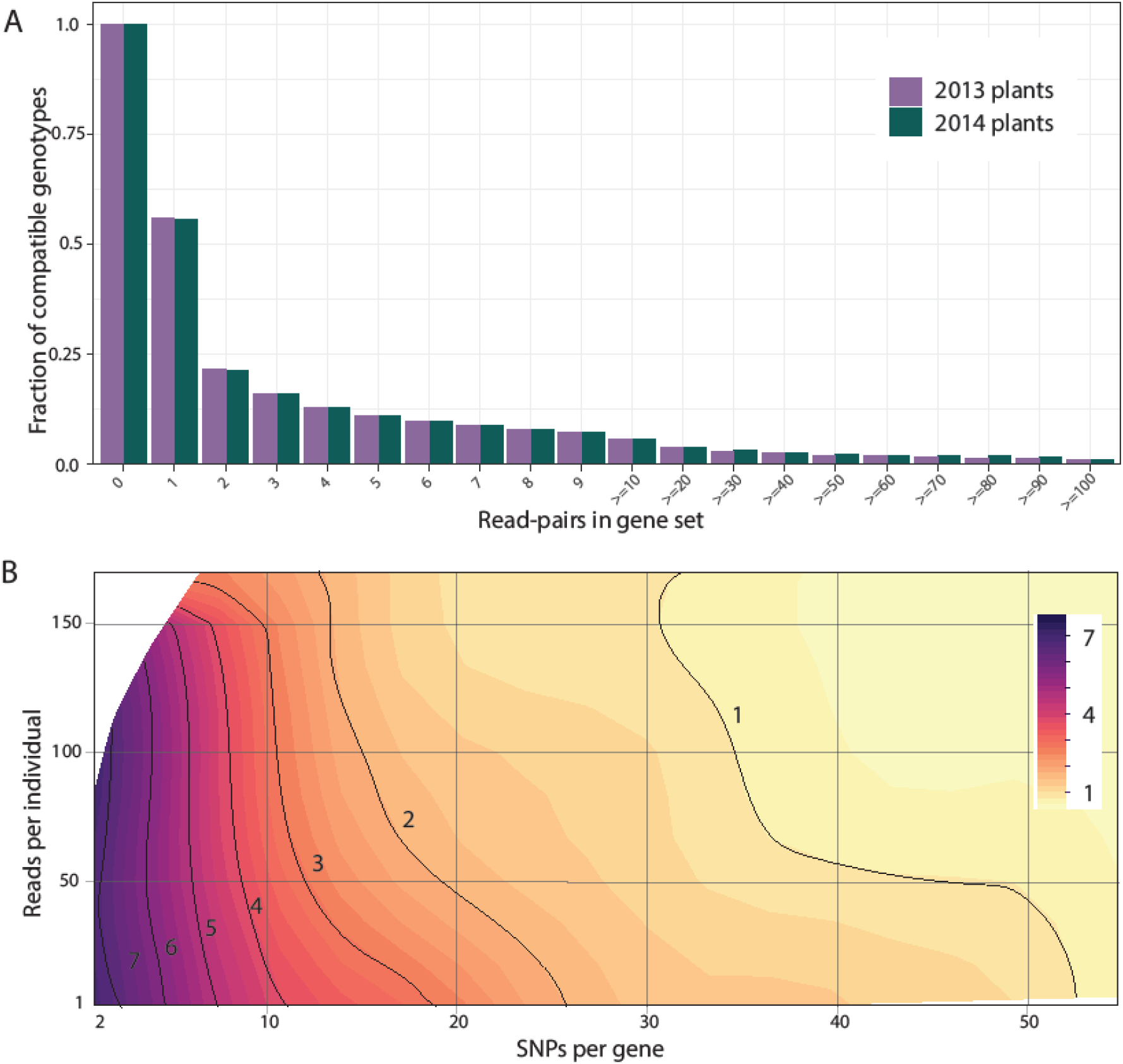
Testing haplotype matching: (A) In Mimulus, the precision of estimation is depicted as a function of the amount of data per plant. Compatible means that the likelihood for a genic-genotype is within 50% of the most likely genotype. (B) In Drosophila, the number of ancestors (indicated by contours and color) matching the genotype of a particular RIL is depicted as a function of amount of data (reads) and the number of SNPs in the gene set.

We collected MSG-RADseq data on 60 of the RILs from DSPR using the same methods as for Mimulus, except that the Drosophila sequences are 94bp single end reads instead of the PE100. We processed the *D. melanogaster* reference genome into ‘gene sets’ and then implemented the same Mimulus pipeline for read mapping, SNP calling and haplotype matching. The great majority of *D. melanogaster* reads overlap 3 or fewer SNPs and are thus less informative than the Mimulus read-pairs (Supplemental Figure S1). Finally, we compared the inferred genotype to the “known” ancestry of each RIL as a test of the method.

This exercise confirms the validity of the haplotype matching, but also its limitations. The ancestral line (or lines) deemed most likely by haplotype matching includes the “correct” line ≈99.5% of the time. We assigned the ancestral genotype as “known” if the posterior probability was greater than 0.99 [42, 51] and thus a small rate of mismatch (less than 1%) is expected even if haplotype matching is perfect. The 99.5% obtained by haplotype matching of MSG data is thus actually close to the theoretical upper limit for accuracy. However, while haplotype matching is accurate, it is not always precise. Oftentimes, the method predicts that numerous genic-genotypes are equally likely. Inference to the specific correct ancestor increases in a predictable fashion with the number of SNPs per gene set and number of reads scored for that line (Fig 2B).

**Figure 3.**
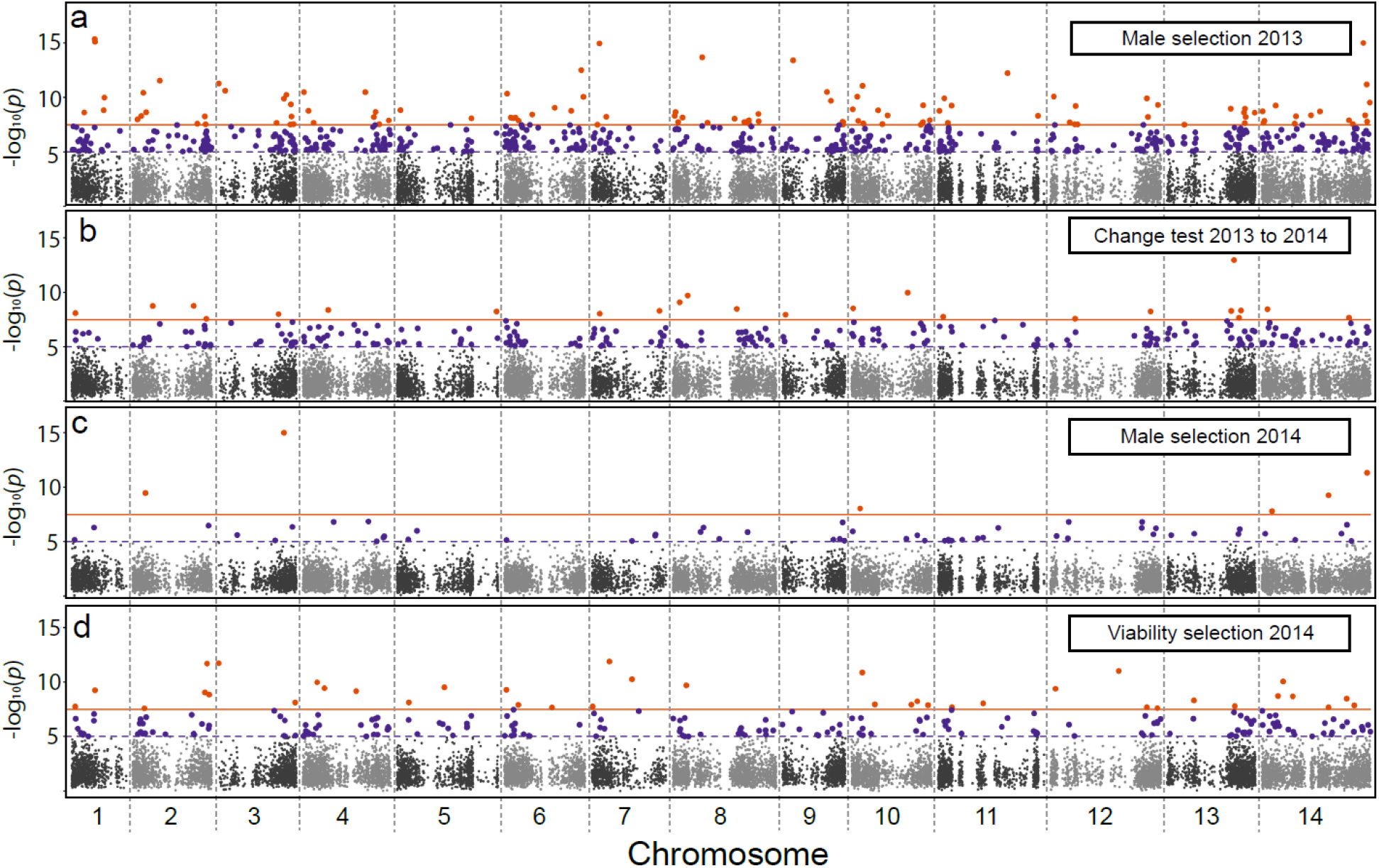
Manhattan plots, with a single test reported per gene, for (a) Male selection 2013, (b) Allele frequency change 2013-2014, (c) Male selection 2014, and (d) Viability selection 2014. The orange line is the Bonferroni threshold, purple is p = 10^−5^.

### Male selection in 2013 predicts change into 2014

We implemented haplotype matching on the Mimulus data and tested 1,523,410 SNPs for selection (filters described in METHODS section B). Testing outcomes within genes were highly correlated owing to linkage disequilibria, and for this reason, we focus on a single test per gene set for the various analyses described below (15,360 tests). Considering the most significant SNP per gene (Supplemental Table S0), 112 tests were genome-wide significant for *p_A_/p_M_* in 2013 (Fig. 3A; Bonferroni α = 0.05/1523410). Given that Bonferroni is excessively conservative, we conducted follow-up analyses accepting SNPs (at most one per gene set) with p < 10^−5^ (587 SNPs in Fig 3A). More false positives (SNPs not under selection) are included with a more permissive threshold, but such SNPs will diminish signal making subsequent tests conservative.

We next performed a test for allele frequency change from 2013 to 2014 by considering the data from both years simultaneously. We first fit a model where *p_A_* in 2013 is constrained to equal *p_Z_*, the allele frequency in zygotes of 2014. We contrast that likelihood to a more general model where *p_Z_* is unconstrained, its value determined entirely by data from 2014. Rejecting *p*_*A*13_ = *p*_*Z*14_ for a SNP indicates allele frequency change into the next generation. Applying this test, we find that 24 genes pass the Bonferroni threshold and that 274 gene sets have at least one SNP with p < 10^−5^ (Fig. 3B). The broad distribution of these tests across chromosomes suggests extensive allele frequency change in IM from 2013 to 2014.

**Figure 4.**
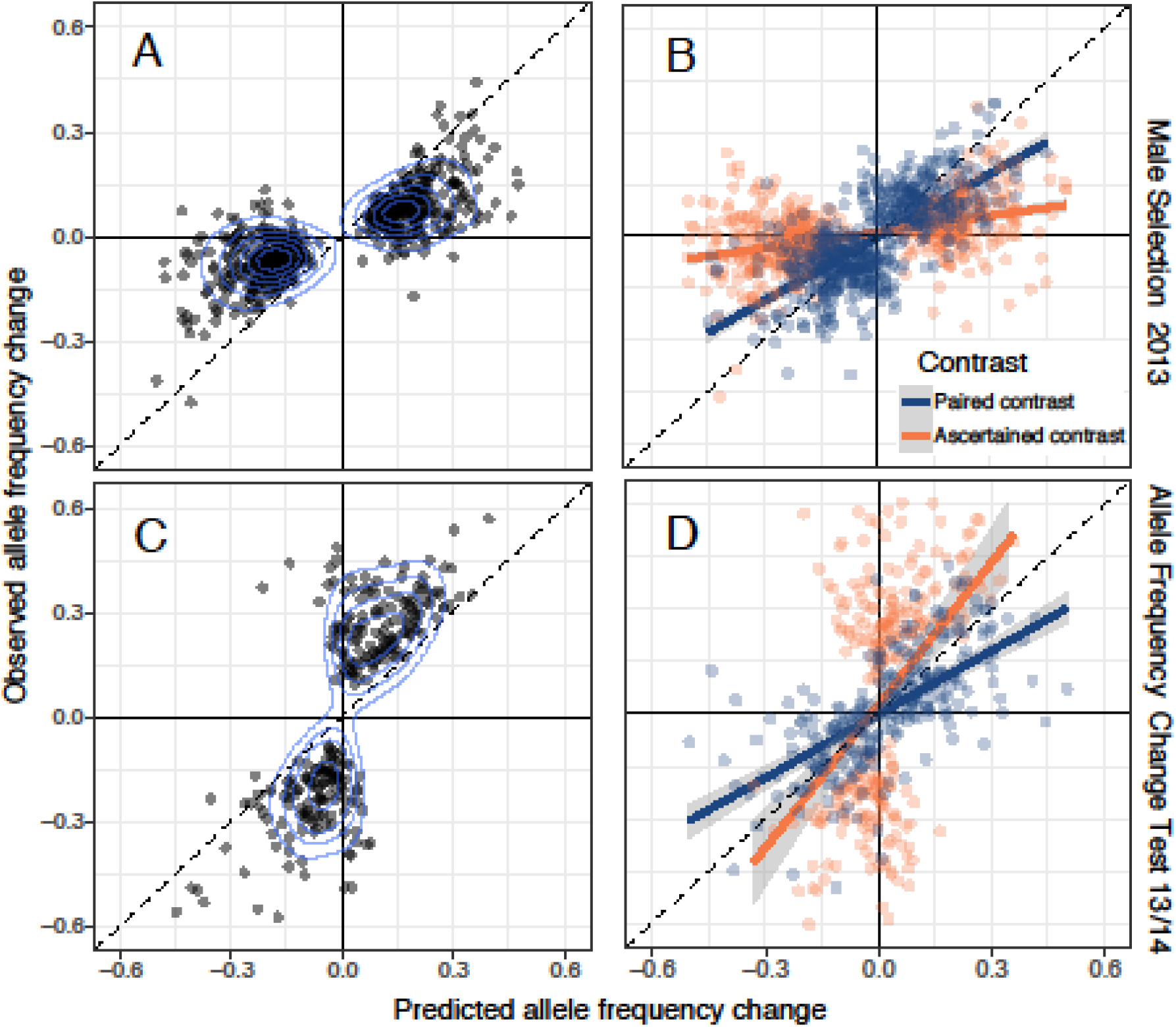
The observed allele frequency change (2013 adults to 2014 zygotes) is compared to predicted with SNPs chosen based on (A) evidence for male selection in 2013 (n = 587) or (C) evidence of change in allele frequency (n=274). Results are reported for all gene sets with a SNP with p < 10^−5^. (B) In the cross-validation of SNPs selected based on male selection, the “Ascertained” contrast is based on the predicted Δp from the significant test (orange points) while the “Paired” contrast is based on the predicted Δp from the other half of the data (blue points). (D) In the cross-validation for allele frequency change significant tests, the ascertained (orange) is the observed Δp from the significant test and predicted Δp from the other data half. Assignment is reversed because the allele frequency change test is based on the observed Δp. For cross-validation, we chose an equivalent number of SNPs to the un-partitioned analyses (n = 587 in (B) to match (A) and n = 274 in (D) to match (C)). Contours indicate the density of points in panels A,C.

We obtain strongly positive relationships between predicted and observed allele frequency change from both the male selection and allele frequency change tests, respectively (Fig 4A: r = 0.79, Fig 4B: r = 0.76, p<0.0001 for both). Both tests imply that alleles elevated in the successful male pollen pool relative to the flowering adult population in 2013 tended to rise in frequency in 2014. We first consider SNPs significant for male selection in 2013 (Fig. 3A) and contrast the predicted change, Δ*p* = (*p_M_* – *p_A_*)/2, to the apparent Δ*p* from 2013 adults to 2014 zygotes (Fig. 4A). Second, we consider SNPs based on evidence for change between years (Fig. 3B) and contrast the direction/magnitude of this observed change to that predicted by 2013 male selection (Fig. 4B). Each relationship deviates from 1:1 (the naïve expectation with unbiased prediction) with the slope for male selection SNPs less than 1 (A: 0.40) and the slope for allele frequency change SNPs greater than 1 (B: 1.57).

The evident positive associations between observed and predicted Δ*p* are very encouraging. However, these relationships require careful statistical scrutiny. The data (and thus estimates) from 2013 and 2014 are statistically independent, but the x- and y-axis Δ*p* values in Fig 4A,C share a parameter (*p_A_* in 2013) that contributes negatively to the Δ*p* estimates on each axis. As a consequence, estimation error in *p_A_* will generate a positive covariance between observed and predicted *apart from that generated by correct prediction*. Ascertainment is second factor. Choosing the most significant SNP for male selection in 2013 will select for those with exaggerated estimates of (*p_M_* – *p_A_*). When male selection favors the reference base, the most significant tests will have positive estimation error added to the true positive value of (*p_M_* – *p_A_*), and the reverse is true for SNPs where the alternative base is favored. The so called “winner’s curse” [52, 53] will thus reduce the regression slope relative to 1 in Fig 4A because the allele frequency in 2014 zygotes is unaffected by estimation error in the previous generation. Ascertainment tends to exaggerate the y-axis variable for the allele frequency change test inflating the slope relative to one. The regression slopes in Fig 4A,C (observed onto predicted) deviate from 1:1 as predicted by this ascertainment effect.

We conducted two analyses that establish genuine prediction of Δ*p* in the face of these errors and biases. First, we used ‘cross-validation’ by splitting the 2013 experiment into odd numbered and even numbered families, respectively. We then performed model fits on each half separately, generating two distinct pairs of observed and predicted Δ*p* for each SNP. We then matched the “odd” predicted Δ*p* to the “even” observed Δ*p*, and vice versa. With this procedure, there is no correlation between observed and predicted in the absence of prediction (Supplemental Appendix D). The partitioning of points in Fig 4B,D reflects the ascertainment step where we choose tests only if male selection (4B) or allele frequency change (4D) was significant. There are two distinct contrasts for a significant SNP. The first is the significant test Δ*p* (say Odd) matched to the observed Δ*p* from the other data half (even), which we denote the “Ascertained contrast.” The remaining data from this SNP (predicted from even, observed from odd in this example) is the “Paired contrast.”

The split data produce strong positive relationships between observed and predicted Δ*p* for both Ascertained and Paired contrasts (Fig 4B,D) despite the reduction in power caused by halving the data. For male selection (Fig 4B), correlations between predicted and observed would be zero for both Paired and Ascertained if SNPs were neutral (or prediction unrelated to response at non-neutral SNPs). In fact, both correlations are highly significant (p < 0.00001 for each in Fig 4B). It is noteworthy that the regression slope is greater for the Paired contrasts (0.62) than the Ascertained contrasts (0.16). This is expected. The magnitude of predicted Δ*p* values is substantially greater in Ascertained relative to Paired contrasts. The exaggeration of predicted Δ*p* inherent to the former group (winner’s curse) reduces the slope. Finally, we note that the predicted Δ*p* is strongly correlated between data halves (r = 0.86, n = 587, p < 0.00001). No correlation is expected under neutrality.

Cross-validation for the allele frequency change test required subdivision of data from both years. We split the 2014 data into even and odd families and (arbitrarily) combined 2013-odd with 2014-odd. Then, as previously, we fit models (here the allele frequency change test) to each data half for each SNP and identified the most significant test per gene. As previously, both Ascertained and Paired contrast sets produce highly significant, positive correlations between observed and predicted Δ*p* values (p < 0.00001 for each in Fig 4D). Here, the regression slope is lower with Paired (0.61) than Ascertained SNPs (1.29). This change in pattern regarding the slopes between in Fig 4B and 4D is predicted given the nature of ascertainment for the allele frequency change test. Here, the observed Δ*p* will be inflated relative the truth for Ascertained but not for Paired contrasts.

As a complement to cross-validation, we developed a full genome simulation program to generate data under the condition that prediction is ineffective (no true relationship between observed and expected). This simulator (Supplemental Appendix D) produces read-pair data equivalent in structure and amount to the real data. To this output, we can apply the full bioinformatic pipeline generating Figs 3–4 from the real data. The simulated data reiterates estimation error and is subject to the same ascertainment biases as the real data, but without real allele frequency change. The latter is assured because we sample genotypes randomly from the set of genic-haplotypes present for each gene set (fitness is equal for all genic-genotypes).

We first applied the selection component models to simulation outputs to confirm our methodology for calling test p-values. We find, that when there is no selection, the sampling distribution of Likelihood Ratio Test values follows the chi-square density (Supplemental Appendix D). This is how we calculated p-values on tests with the reals data. The observation of chi-square distributed LRT values simulations confirms the asymptotic normal theory for likelihood testing. Second, we confirmed that the cross-validation method eliminates the spurious association between predicted and observed Δ*p* (null hypothesis for Figs 4B,D). Finally, the simulations confirm that a positive association between observed and predicted change is generated by estimation error in the un-partitioned data (Fig 4A,C). However, the covariance between observed and predicted is much greater for the real data than for the simulated data (0.020 vs 0.012 for male selection, 0.033 vs 0.012 for the allele frequency change test). Thus, the magnitude (if not simply the direction) of the covariance in Figs 3A,C is indicative of effective prediction.

In summary, the simulation and cross-validation procedures provide strong support that prediction is genuine. Unfortunately, it is much more difficult to determine the extent that apparent deviations between observed and predicted are due to sampling error as opposed to model error. The regressions of observed onto predicted Δ*p* for Paired contrasts (Fig 4B,D) are the simplest parametric relationship to interpret. The slopes for these, 0.61 and 0.62, suggest that response is less than predicted, but this conclusion is very tentative. Simple estimation error in the predictor of a linear regression causes a downward bias in the slope (here relative to one), even when there is no ascertainment bias [54]. This is non-trivial given that our SNP-specific predictions (and observations) of allele frequency change are encumbered with substantial estimation error. The relationship between estimation precision and experimental design, including sample size, is demonstrated in the companion paper [50].

Several biological factors may have reduced model accuracy. For example, we assumed that (a) there was no differential germination in the greenhouse (affected by genotype) when we grew progeny from maternal plants of 2013, (b) no seed bank contributed to the 2014 generation, and (c) no immigrant pollen or seed contributed to the 2014 population. Each of these influences could cause systematic deviations between observed and predicted Δ*p*. Germination rates routinely differ between plant genotypes in an environment-dependent fashion, e.g. [55, 56]. The field environment of 2014 (where plants germinated to produce our observed Δ*p*) is certainly different from the greenhouse (the offspring genotypes used to estimate *p_M_* in 2013). This could cause substantial deviations, although they would be limited to genomic regions containing “germination genes.”

Prediction accuracy for many loci could be affected by the violations of the other assumptions: (b) seed bank or (c) gene flow. If selection varies substantially among years, and all evidence indicates that IM experiences strong fluctuations ([17, 22–25, 27] and results below), a seed bank can moderate temporal changes in allele frequency [57]. *M. guttatus* does not have seed dormancy [58], and at present, we have no evidence that a seed bank exists for IM. If it does however, recruitment from the seed bank would probably act to reduce the magnitude of observed Δ*p* relative to predicted. Finally, there certainly is some level of gene flow into IM from other populations [49]. However, the fact that IM is a very large population [49], coupled with the observation of substantial allele frequency divergence from neighboring population [59], suggest that the rate of immigration is quite low (<< 1%). This level of gene flow might fundamentally alter long-term evolutionary dynamics (e.g. by introducing novel alleles), but should not have a dramatic effect on single-generation Δ*p* values.

### Regularities in genome-wide selection

In the previous section, we used the 2014 data simply to estimate the observed Δ*p* from selection in 2013. However, the experimental design for 2014 allows a more detailed dissection of fitness variation within this generation. Viability selection estimated from the difference between *p_A_* and *p_L_* (Fig 1B) was abundant: 39 genes pass the Bonferroni threshold and 226 have at least one SNP tests with p < 10^−5^ (Fig 3D). Male selection was considerably weaker in 2014 than 2013: only 6 *p_A_/p_M_* tests pass Bonferroni, 59 genes have a SNP with p < 10^−5^ (Fig 3C). The pattern of selection also changed. In 2013, there was a clear tendency for male selection to favor the minor (less frequent) allele (Fig 5A). The average predicted change of the minor allele frequency (MAF) was 0.055 (SE=0.009), which is significantly positive (n = 587, t = 6.16, p<0.001). In 2014, the predicted change in minor allele frequency caused by male selection was close to zero. These data corroborate previous studies demonstrating changes in the direction/magnitude of genome-wide selection between generations, both in Mimulus [17] and other systems, e.g. [12]. Absent such fluctuations (or other trade-offs), we would expect rapid fixation of one allele or the other, and the loss of fitness variation.

**Figure 5.**
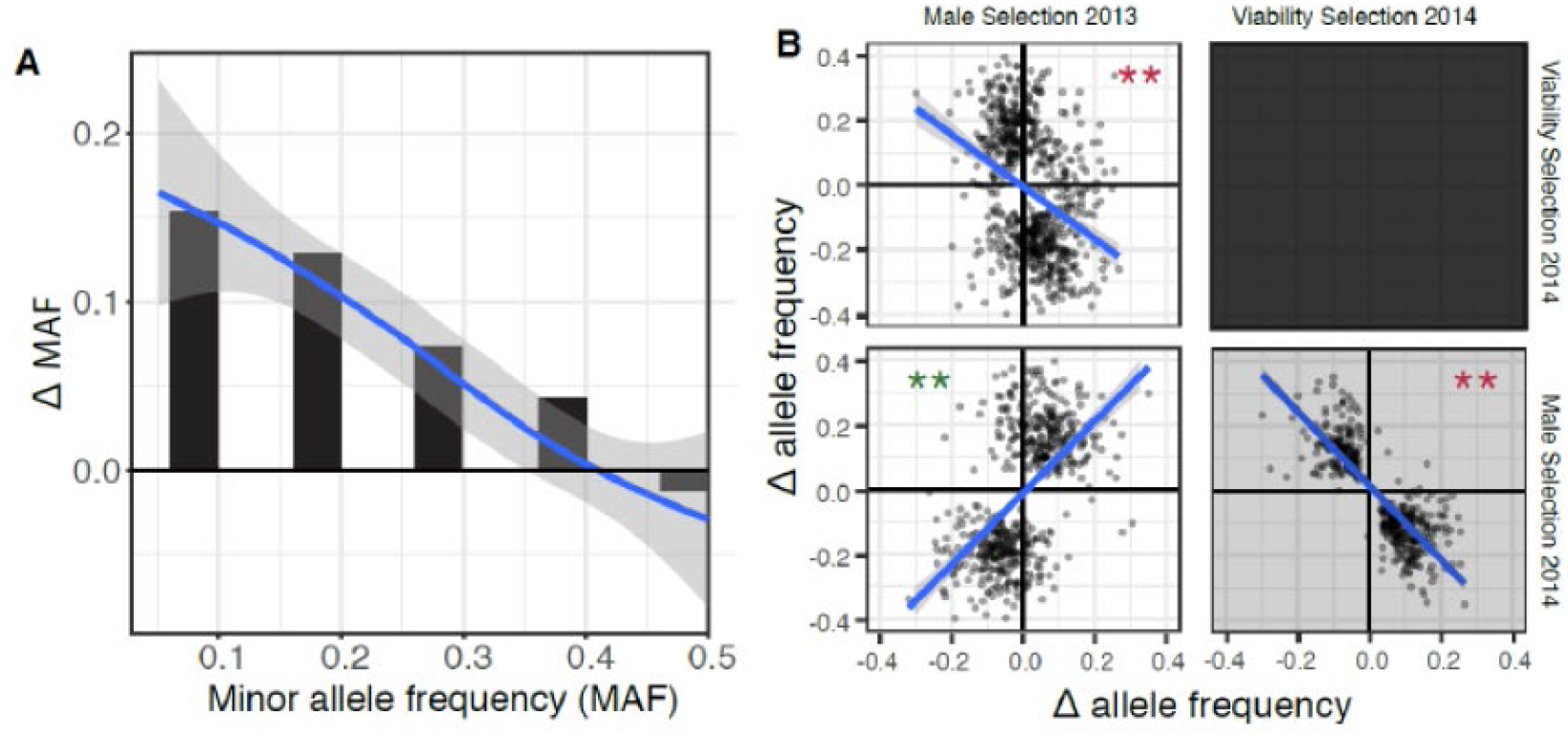
(A) Male selection favored minor alleles in 2013. (B) Pairwise contrasts between predicted changes owing to male selection in 2013, viability selection in 2014, and male selection in 2014. A single SNP per gene is reported (the most significant) if p < 10^−5^.

Selection components exhibit strong correlations indicating consistency in male selection across years and a trade-off between male selection in 2013 and viability in 2014 (Fig. 5B). To compare different components of selection, we selected the SNP within each gene set with the highest aggregate evidence for selection using Fisher’s combined probability statistic [60]. Alleles favored by male selection in 2013 were also favored by male selection in 2014 (n = 555, r = 0.57, p<10^−48^), but disfavored by viability selection in 2014 (n=725, r = −0.34, p<10^−20^). As expected from these results, there is also a negative correlation between male selection and viability within 2014 (r = −0.83), but testing is complicated for this contrast because the two tests share a common parameter and thus subject to biases discussed previously.

We can perform a final contrast with a previous experiment that transplanted IM genotypes as seedlings into a neighboring field site at Browder Ridge. Browder Ridge has similar physical conditions as IM [17, 24, 25, 27]. The 2014 transplant [17] assayed 62 distinct IM genotypes for survival, each a cross between two of the 187 sequenced IM lines. Of SNPs indicating viability selection here (p<10^−5^; Fig 3D), 28 showed at least suggestive evidence of viability selection in the transplant experiment (test p-value less than 0.1). Despite the small sample size, there is a strong positive correlation between predicted Δ*p* between the two independent experiments (r = 0.53, p < 0.004).

The contrast across years (Fig 5B) is an important confirmation of natural selection as the principle driver of Δ*p*. If apparent changes were caused entirely by sampling and/or estimation error, the direction of change would not be correlated between independent datasets (2013 versus 2014 plants). Recent studies in fully pedigreed populations of birds and mammals have clearly shown substantial allele frequency change through time [21, 61, 62]. The challenge has been to attribute changes to natural selection as opposed to genetic drift [21, 61]. In the present study, the sampled population (n is about 1000 individuals from each year) is orders of magnitude smaller than the number of reproductive individuals within the population each generation [49]. The null hypothesis in our tests for selection is essentially experiment-level drift (differences in allele frequency caused by the finite numbers of parents and offspring). Experiment-level drift is necessarily much stronger than population level drift because n << N. Significant tests thus clearly implying selection, albeit with the caution that negative results (non-significant tests) do not imply that SNPs are evolving neutrally. Thus, undetected selection through measured components, as well as selection via unmeasured fitness components make our results a conservative picture of the genome-wide extent of natural selection.

### Conclusions

This experiment demonstrates strong, but often antagonistic, selection on hundreds of genes (Figs 3–5). The apparent trade-off between fitness components, as well as the correlations between allele frequency and direction of Δ*p*, extend and corroborate previous experiments on this population. Figure 5B provides further evidence that Montane annual populations of *M. guttatus* exhibit a life-history trade-off between development rate and reproductive capacity. In most years (although not 2013 of this experiment), nearly all plants die owing to drought at approximately the same time, but *survival to flowering* differs greatly owing to varying rates of maturation [27, 63]. The current study shows clear evidence of a viability trade-off with male reproductive success, with male selection for minor alleles in 2013 likely mediated through positive effects on flower size in this year of favorable growth conditions. Furthermore, consistency between 2013 and 2014 in the direction of allelic effects on male fitness suggests that such tradeoffs are intrinsic and contribute to the maintenance of big/slow alleles at minor frequencies within IM [17, 26]. This is yet another of a growing body of examples relating antagonistic pleiotropy to polymorphism across diverse systems, e.g. [64].

Selection on both quantitative traits and specific genetic loci with major effects can be quite strong [1, 2, 65]. However, both conceptual and logistical difficulties have separated phenotype-level and locus-specific approaches, limiting inference about the extent, nature, and magnitude of selection on genetic variants across the genome. Our results (Fig 3) suggest that genotypic fitness is broadly estimable, and that these estimates can predict allele frequency change across generations (Fig 4). A shortcoming of this study (considered in isolation from prior work at IM) is that the selection component estimates do not provide an ecological explanation for the observed selection on SNPs. As in quantitative genetics, we can obtain such an understanding by replicating the measurement of selection across different populations (or the same population through time) and then correlating selection estimates with environmental or ecological variables. Mechanistic insights may also come from combining phenotypic measurements with genotyping and fitness assays, linking GWA with selection component analyses. In summary, a broader application of genomic selection component methods, coupled with environmental/phenotypic data and population monitoring through time, should help to resolve the limits of population genetic prediction.

## Materials and methods

### A. Field sampling and progeny testing

*Mimulus guttatus* (syn *Erythranthe guttata*) is a wild flower species (Family: Phrymaceae) abundant throughout western North America [66]. The IM population, located in the central Oregon cascades (44.402217 N, −122.153317 W, Elevation ~1400 meters), is described in detail elsewhere [22, 24, 27]. In 2013, whole plants distributed in a grid across the IM population were collected (at senescence) into coin envelopes. In 2014, we established three primary transects (each ~10m) horizontally across the face of the slope, with approximately equal vertical spacing between transects. The transects were further subdivided into perpendicular sub-transects which extended 0.3m on either side of the primary transect and were evenly spaced in 0.3m increments along the primary transect. We sampled five plants along each sub-transect by selecting the most proximal individual to a points placed at 10cm intervals. On July 15, 2014, we surveyed each transect and identified plants that would not progress to flower based on state of development relative to others in population. Assuming these plants would not have sufficient time to flower and set seed prior to season ending drought, this cohort (L) estimates *p_L_* in Fig 1. To insure sufficient DNA from L plants, we transplanted these individuals into moistened peat pots filled with potting soil and reared them to sufficient size for DNA extraction. We first sampled plants for the adult cohort of 2014 (*p_A_* in Figure 1) on July 21, 2014. We only sampled adults once all plants within their sub-transect fully dried down. We collected whole plants, after confirming they had begun setting seed, into envelopes, so that both seed and maternal tissue could be separated for planting and DNA extraction, respectively. The remaining adults were harvested on July 27. Given seed collections from both years, we germinated and grew 2-4 progeny from each field plant in the University of Kansas greenhouse. We harvested dried leaf and calyx tissue from field collected parental plants and young leaves from greenhouse germinated progeny for subsequent DNA extraction[67]. To determine the overall proportion of the population that survived to flower in 2014, we surveyed a random set of 1000 seedlings marked early in the season at the nearby BR location[59]. 700 of these plants eventually flowered.

### B. Library preparation, sequencing, SNP calling, and scoring read pairs

We collected paired-end sequence reads from 1936 experimental plants (2013: 207 field plants and 685 progeny; 2014: 383 field plants and 661 progeny) using Illumina technology. For field plants and their progeny, we generated genomic libraries using Multiplexed-Shotgun-Genotyping (MSG)[41], a form of RADseq [68] that uses a restriction enzyme to reduce genomic representation to homologous loci that are flanked by restriction cut sites. We digested genomic DNA from each plant using the frequent-cutting restriction enzyme MseI (NEB Biolabs). Each DNA sample was ligated to one of 96 distinct barcoded adaptors, each containing a unique 6 bp barcode. Each set of these barcoded samples is then pooled independently to create a sub-library. After PCR, we size-selected our library for 250-300bp fragments using a Pippin Prep (http://www.sagescience.com/products/pippin-prep/). We then performed PCR reactions at 12 cycles using Phusion High-Fidelity PCR Master Mix (NEB Biolabs) and primers that bind to common regions in the adaptors. In the PCR step, each sub-library was combined with one of 24 distinct Illumina indices allowing multi-plexing of the sub-libraries. To remove primer dimers, we did two rounds of AMPure XP bead cleanup (Beckman Coulter, Inc) using a 0.8 bead volume to sample ratio. Libraries were sequenced with 100-bp paired-end reads on the Illumina HiSeq 2500 with a 10% phiX spike-in. The specific program commands used to call SNPs in the MSG data are described in Supplemental Appendix A. We suppressed Indels and all SNPs with more than two nucleotides segregating.

Sequencing and variant calling on the 187 reference panel genomes from IM was described previously[17]. We first imputed the few missing calls in these genomes and then extracted the sequence for each reference genome within each gene set (detailed procedures in Supplemental Appendix B). Sequence variation is very high in *M. guttatus[49*], and as a consequence, it is difficult to effectively call variants outside genic regions. We thus established gene sets as units for analysis. A set is either a single gene or a collection of closely linked (within 100bp) and/or overlapping genes. After suppressing genes prone to paralogous or otherwise spurious read mapping, 15,360 gene sets were retained for subsequent analysis (Supplemental Table S1). Finally, we noted that some SNPs were completely redundant – owing to perfect association in the reference panel, they always produced the same genotype likelihoods in field plants. We thinned sets of fully redundant SNPs to a single representative SNP leaving 1,523,410 SNPs for selection estimation.

The data units for likelihood calculations (eq 1) are read-pairs scored for each polymorphic SNP that they overlap within a gene set. We aligned the read-pairs from each plant to the whole genome sequences, and within each gene set, and calculated *U*_[*plantID*]*i,j*_ for each possible genic-genotype [i,j]. *U*_[*plantID*],*i,j*_ is the likelihood for the full collection of read-pairs from a plant given that its diploid genic-genotype is [*i,j*], where *i* and *j* index genic haplotypes. Based on the low mismatch rate to genic haplotypes (as a whole), we set *ϵ* = 0.005 for calculation of eq (1) described below. We calculated *U*_[*plantID*],*i,j*_ for each combination of gene set, plant, and genic-genotype using python scripts p1.py, p2.py, p3.py, p.Uij.2013.py and p.Uij.2014.py (Supplemental file 1). Application of these programs indicate that some closely linked SNPs were completely redundant – they always had exactly the same genotype calls in field plants. We thinned these cases to a single SNP.

### C. Drosophila melanogaster analysis

The *Drosophila* Synthetic Population Resource (DSPR) consists of two multiparental, advanced generation intercross mapping populations [42, 51]. Each population (A and B) was initiated with eight inbred founder strains, with one strain common to both populations (i.e., 15 founders in total). Following 50 generations of free recombination, a series of Recombinant Inbred Lines (RILs) were initiated by 25 generations of sibling mating. The founder genomes were sequenced to 50X coverage and the RILs subjected to RAD-seq using SgrAI, an 8-cutter, as the restriction enzyme [42, 69]. Given these data, we are able to infer the mosaic founder haplotype structure of each RIL at >10,000 positions covering the genome.

We collected MSG RADseq data using the same protocol as described above for the Mimulus experiment, except that these data are 94bp single end sequences instead of the PE100 sequencing for Mimulus. We chose 60 of the RILs for the present study equally split between set A and set B of the DSPR. For each collection, there are only 8 possible ancestral genomes, but we ran the analysis blind to this information (thus inference among 15 possible ancestral alleles was required). The reads were processed with fastp (https://github.com/OpenGene/fastp) and then we mapped to the FlyBase r5.56 genome build (https://flybase.org/) and called SNPs following the procedures used for Mimulus (Supplemental Appendix A). We used the annotation (dmel-all-r5.56.gff) to establish a list of 13,384 gene sets applying the same rules as for Mimulus (Supplemental Table S3). Next, we determined the intersection between SNPs within the ancestral genomes (final_snptable_foundersonly.txt downloaded from http://wfitch.bio.uci.edu/~dspr/) and those called in the MSG RIL data, a total of 107,878 bi-allelic SNPs (Supplemental Table S4). We found that 8900 of these 13,384 gene sets had at least one SNP scored in MSG data and could thus be used for downstream analysis. After eliminating uninformative reads, a total of 15,488,651 remained across the 60 RILs. We next adapted the Mimulus programs (python scripts p1.py, p2.py and p3.py in Supplemental file 1) to determine predicted ancestry based of the DSPR RILs and matched the inferred ancestry to the “known” ancestry of each RIL. The latter was established previously: We downloaded files HMMregA_R2.txt and HMMregB_R2.txt from http://wfitch.bio.uci.edu/~dspr/ (also available at https://datadryad.org/stash/dataset/doi:10.5061/dryad.r5v40). We processed the *D. melanogaster* reference into ‘gene units’ by the same method applied to the Mimulus genome. Read mapping and SNP calling were executed using the same techniques. The great majority of *D. melanogaster* reads overlap 3 or fewer SNPs and are thus less informative than the Mimulus read-pairs (Supplemental Figure S1). We then applied the inference programs using the 15 ancestral sequences of the DSPR as genic haplotypes.

### D. Likelihood of the field data with and without selection

Selection component analyses (SCA [43, 46]) are based on population genetic models that predict allele frequency change from observations of viability, fecundity, and mating success [47]. SCA estimate selection from differences in allele frequency between distinct “cohorts” within a population, e.g. individuals that survive to reproduce and those that do not (viability selection) or those that acquire mates and those that do not (sexual selection) [48]. Given random sampling of individuals, the likelihood of the entire dataset (*L*) is a product across families:

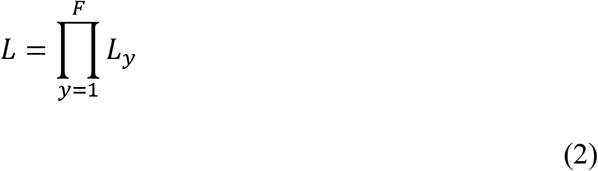

where *F* is the number of families and *L_y_* is the likelihood for family *y*. Families consist of a single individual if that plant failed to survive to reproduce. For survivors, the family is the field plant and a sample of their progeny. The log-transformed likelihood:

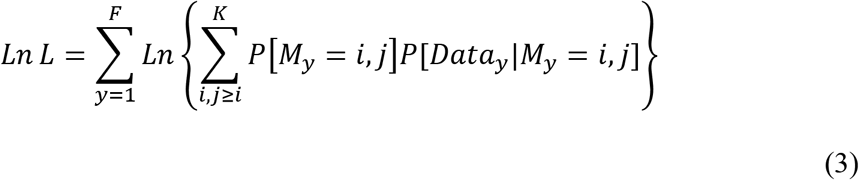

where *P*[*M_y_* = *i, j*] is the (prior) probability that the maternal genic-genotype has genic-haplotypes *i* and *j*. *K* is the number of distinct sequences for this gene set. *P*[*Data_y_*|*M_y_* = *i, j*] is the probability of all data from family *y* (genetic and fitness measurements) given maternal genotype [*i,j*]. The family likelihood is:

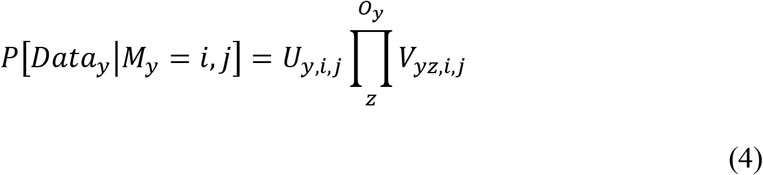

*U_y,i,j_* is the probability maternal plant *y* produced the observed read-pairs given genic-genotype [*i,j*], *V_yz,i,j_* is the probability of the observed read-pairs for offspring *Z* of maternal plant *y* with genic-genotype [*i,j*], and *O_y_* is number of genotyped offspring of maternal plant *y*. For individuals that fail to reproduce, [*Data_y_*|*M_y_* = *i, j*] = *U_y,i,j_*. The likelihood for each offspring, *V_yz,i,j_* in eq 4, depends on whether that offspring is outcrossed or selfed (see Methods section E). If offspring *yz* is selfed:

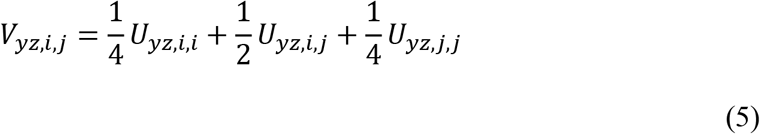

We assume that each outcrossed progeny is sired independently and that

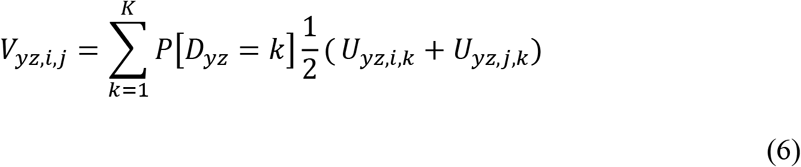

*U_yz,v,w_* is the probability of the observed read-pairs from offspring *yz* given that it has genic-genotype [v,w]. *P*[*D_yz_* = *k*] is the probability that the sire of offspring *yz* transmitted genic-haplotype k to this offspring. The (1/2) reflects the equal probability of transmission for either maternal allele (*i* or *j*) to the offspring. Through all these calculations, we assume that recombination within gene sets has a negligible effect on the probabilities.

The various models of selection (Fig. 1) consider different constraints on the genotype probabilities. Given the large number of genic-genotypes, the potential parameter space is very large. Here, we simplify by classifying all genic-haplotypes into two groups based on their allele at a particular SNP. We assume the sequences in a group are equivalent in terms of fitness effects. This reduces all genic-haplotypes at a gene set into two “alleles” for selection tests. This classification naturally changes with SNP chosen and thus we apply the procedure to each SNP in sequence. This simplification is a sensible first step, but we acknowledge that it may fail to capture the genotype-to-fitness mapping for many genes. In some cases, alternative alleles may be defined by numerous SNPs or indels within a gene [70, 71] and fitness effects would be more naturally described with an allelic series. Our ‘binning’ of functionally distinct alleles could elevate the Type I error rate (we fail to see selection when it is occurring).

Let *S_R_* represent the set of genic haplotypes that have the reference base at the focal SNP and *S_A_* is the set with the alternative base. Then eq (6) can be written:

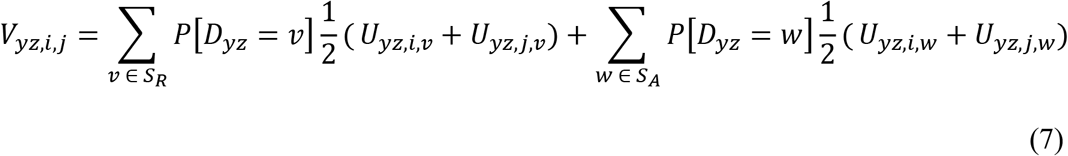

The frequency of the reference base (for the focal SNP) within the population of genic-haplotypes, *p*, is just 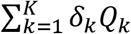, where *Q_k_* is the frequency of haplotype k among the lines and *δ_k_* is an indicator variable (1 if haplotype *k* carries the reference base and 0 otherwise). Of course, the frequency of the reference base can differ between the sequence line set and the natural population, and also between subsets of the natural population (e.g. alive versus dead). Let *p*\* denote the frequency of the reference base in a specific field cohort, say adults in 2013 or zygotes in 2014. We adjust genic-haplotypes proportionally as a function of *p*\*:

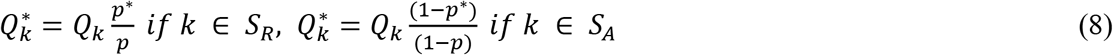

This is essentially a uniform inflation or deflation of haplotype frequencies based on the focal SNP. It allows us to write the likelihood equations explicitly in terms of allele frequencies at one SNP (e.g. *p_L_*, *p_A_*, and *p_M_* in Figure 1) while retaining the full information from gene sets. For example, *P*[*M_y_* = *i,j*], in eq (3) becomes 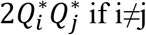 or 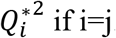. This is a function of known fixed values (p, *Q_i_*, *Q_j_*) and the parameter to be estimated (e.g. *p_A_* if the maternal plant survived, *p_L_* if not). Equation (7) becomes:

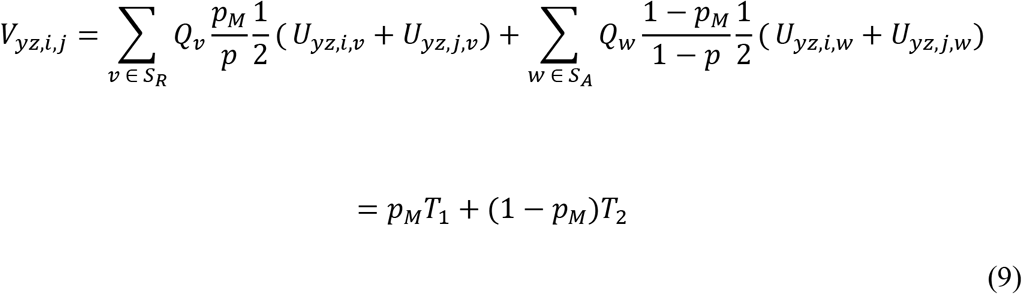

*T*_1_ and *T*_2_ distill all quantities in eq (9) that are coefficients for *p_M_* and (1 – *p_M_*). The fact that these coefficients are determined entirely by the read-pairs from field plants and the set of genic-haplotypes means that they do not change with *p_M_*. Thus, the numerically intensive sum of eq (6) need only be calculated once at the onset of a maximum likelihood search. We use Powell’s algorithm [72] to maximize likelihoods. At each SNP, we fit a series of models of increasing complexity (Fig. 1). Likelihood ratio tests are used to evaluate whether more general models are superior to simpler models. The code to perform these tests was written in the C programming language, is described in Supplemental Appendix C, and is included in Supplemental File 1.

### E. Mating system estimation

The MSG data (without the reference sequences) was used to determine individual offspring as outcrossed or selfed using BORICE [73]. The most informative SNPs for mating system estimation exhibit high coverage across samples and intermediate allele frequency. From the full set of MSG samples called simultaneously (Supplemental Table S5), we chose one SNP per gene with the highest count for (heterozygotes+the less frequent homozygote) using python program p4.py (Supplemental File 1). We then extracted genotype likelihoods for these SNPs (directly from the vcf file, Supplemental Table S5) and organized the samples into families (maternal plants with offspring) to produce a BORICE-format input file using python program p5.py (Supplemental File 1). We next thinned the dataset to SNPs with at least 800 called plants (across both years) producing the input file used for estimation of mating system (Supplemental Table S6) consisting of 2773 SNPs, each in a distinct gene. These SNPs are well distributed across all 14 chromosomes. We conducted preliminary MCMC runs to determine parameter step sizes, burn-in duration, and chain length. After setting these (Control file and the specific BORICE code are in Supplemental File 1), we estimated posterior probabilities for each offspring as outcrossed/selfed and the inbreeding level of maternal plants by combining four independent chains.

Considering offspring with at least one read at 100 or more SNPs, 10.1% were determined to be selfed in 2013 (54 of 537) versus 9.4% in 2014 (48 of 508). The remaining offspring, where there was insufficient data for estimation, were set as outcrossed for the subsequent selection analyses. While this classification may be incorrect for a few individuals, error has a minimal effect on parameter estimates given the absence of genotypic data for these offspring. The observed rate of selfing (ca. 10%) matches results from prior mating system studies of the IM population[74]. The detailed results are reported in Supplemental Table S7.

### F. Predicted and observed allele frequency change

We contrast different selection estimates in the common currency of predicted allele frequency change, Δ*p*. Considering the change from adults to zygotes of the next generation, the predicted change due to male selection is Δ*p* = (*p_M_* – *p_A_*)/2. This equation assumes no differential female fecundity (associated with the SNP) and that all progeny are produced by outcrossing (diploid loci are half male and half female). In fact, we found that ca. 10% of our offspring were derived from selfing (see section **E**). This could (slightly) inflate predicted change relative to observed change (Fig 3). However, given that the inflation is uniform, it does not affect arguments about significance (Fig 2), allele frequency (Fig 4) or trade-offs (Fig 4). The predicted change owing to viability selection in 2014 is calculated from model H_3_ (Fig 1) estimates, *p_A_* and *p_L_*. The relevant relationship is *p_Z_* = *α p_A_* + (1 – *α*) *p_L_*, where *p_Z_* is allele frequency in zygotes (before selection) and *α* is the fraction of individuals that survive to reproduce. For our experiment, we estimate *α* = 0.7 (see above in section **A**). Rearranging the equation, the predicted change owing to viability selection is Δ*p* = 0.3(*p_A_* – *p_L_*). The observed Δ*p* estimates (Fig 3) require an estimate of allele frequency in zygotes (*p_Z_*) from 2014. This can be estimated in several ways given the four models applied to the 2014 data (H_0_-H_3_ in Fig. 1), but *p* from H_0_ is a robust choice. This value is always intermediate to parameter estimates from models that are more elaborate.

To obtain Δ*p* from the viability data from the transplant experiment in 2014 at BR[17], we first determined the 1,358,005 SNPs in common between the SCA (results of this study) and the genotypes used in that study. We assayed 355 transplants (an average of 5.7 replicates per genotypes) for survival and seed set of survivors in the 2014 transplant. We eliminated SNPs where the count of minor homozygote plus heterozygotes was fewer than five. For the remainder, we regressed the fitness measure, either fraction surviving or mean Ln(seedset of survivors), onto plant genotype at each SNP, the latter scored as the count of Reference alleles (0,1,2). A linear model for selection was used instead of estimating the mean for each genotype (RR, RA, AA) because there were often few or no representatives of the minor homozygote (only 62 distinct hybrid genotypes were assayed in the 2014 transplant). For viability selection, the predicted change is Δ*p* = *b_v_p_Z_*(1 – *p_Z_*)/ *w_v_*, where *b_v_* is the regression coefficient and *w_v_*=0.46 is the mean viability among transplants. Allele frequency, *p_Z_*, was taken from the 2014 SCA model H_0_.

## Supporting information

Supplemental Appendix. The detailed methods sections for (A) Bioinformatic processing of MSG data, (B) Delineating gene sets and SNPs, (C) Selection component models, and (D) Whole-genome data simulation.

Supplemental Table S0. The most significant SNP per gene is reported for *p_A_/p_M_* in 2013, *p_A_/p_M_* in 2014, viability selection in 2014, and the change test (2013 adults to 2014 zygotes). The chosen for each test are reported on a separate sheet. Statistics from all model fits are reported for each SNP.

Supplemental Table S1. The gene sets are located to the genome sequence and the number distinct genic-haplotypes per gene set is reported.

Supplemental Table S2. The number of SNPs covered per read-pair in the Mimulus field plants. After discarding read-pairs that overlap no SNPs, slightly more than 99 million remained.

Supplemental Table S3. The collection of genes and gene sets for the Drosophila application: “Gene.coordinates.txt”.

Supplemental Table S4. Variants used in Drosophila application: “SNPs.in.both.txt”

Supplemental Table S5. The Variant Call File (vcf) for all msg samples across both field seasons is given for each chromosome separately.

Supplemental Table S6. The BORICE formatted input file for mating system estimation.

Supplemental Table S7. The estimated posterior probabilities that each offspring is outcrossed and for the Inbreeding History (IH) level of maternal plant is reported.

Supplemental Figure S1. The number of SNPs per read (Blue = Drosophila) or read-pair (Orange = Mimulus) is reported as a histogram.

File S1 key. A key to the programs contained in Supplemental File 1.

Supplemental File 1. The 14 programs used to analyze and simulate data (detailed descriptions contained in File S1 key).

## Acknowledgements

We thank C. Friesen (U.S. Forest Service) for site access and the KU ACF for computing resources. J Stinchombe suggested the data splitting for cross-validation and we received essential editorial advice from J. Willis and R. Unckless. Sequencing was conducted at the KU genomics core (supported by the CMADP COBRE P20GM103638).

## Author Contributions

PJM and JKK conceived the project. PJM and LF conducted the field experiments. PJM directed library construction and sequencing for Mimulus. SJM directed library construction and sequencing for Drosophila. JKK wrote the theory and the analytical programs. JC and JKK analyzed the data. JKK wrote the paper with substantial input from all co-authors.

## Availability of data and materials

The Illumina reads from both Mimulus and Drosophila will be deposited in the Sequence Read Archive (NCBI) prior to publication. Computer programs to conduct the analyses have been included as Supplementary Materials.

